# Association of human gut microbiota with Alzheimer’s disease pathogenesis: An exploratory clinical study

**DOI:** 10.1101/2025.01.28.635229

**Authors:** Tadashi Ohara, Yasuyuki Taki, Shinya Yamamoto

## Abstract

The relationship between Alzheimer’s disease (AD) onset and the brain–gut axis has garnered increasing attention. This study aimed to investigate the potential role of the brain–gut axis in AD pathogenesis, with a specific focus on microbiota composition. This exploratory study enrolled 10 patients with AD and 13 healthy adults, grouped by age (≤30 years, 31–40 years, and ≥41 years). Fecal samples were collected, and 16S rRNA gene sequencing was employed to analyze differences in fecal microbiota composition at the bacterial species level. Certain bacterial species appeared more abundant in the AD group (e.g., *Ruminococcus inulinivorans* and *Ruminococcus torques*), while others were relatively more abundant in healthy adults (e.g., *Prevotella vulgatus 1*, *Bacteroides wexlerae*, *Clostridium butyricum*, and *Alistipes rectalis*). However, these differences were not statistically significant, likely because of the limited sample size. These findings suggest that fecal microbiota composition may differ between patients with AD and healthy individuals, with a potential intermediate group at risk of AD development. Larger-scale clinical studies are necessary to further elucidate the bacterial species associated with AD pathogenesis, potentially enabling the use of microbiota composition as a screening tool to distinguish between healthy individuals, patients with AD, and those with preclinical AD.

## Introduction

The increasing prevalence of dementia poses a significant societal challenge. Alzheimer’s disease (AD), which accounts for approximately 70% of all dementia cases, progresses slowly and is characterized by the accumulation of amyloid-β (Aβ) peptide in the brain and the hyperphosphorylation of tau protein, resulting in brain tissue damage [1–3]. Among Aβ peptides, Aβ42 is highly hydrophobic and amyloidogenic, and elevated levels of Aβ42 are strongly associated with amyloid plaque formation [4,5,6]. Interestingly, studies have shown that germ-free mice lacking intestinal microbiota do not exhibit Aβ accumulation in the brain [7].

Recently, it has been reported that transplantation of human feces from patients with AD into healthy rats causes AD in these rats [8] and that transplantation of feces from young mice into aged mice results in the improvement of cognitive function and immunity [9]. Of particular interest is the intestinal bacterium *Clostridiales*, which produces an Aβ-like peptide with high homology to the human Aβ peptide, sharing a similar tertiary structure and Aβ42-like activity [10]. Experimental evidence from various mouse models indicates that Aβ peptides produced by intestinal bacteria can enter the bloodstream, cross the blood–brain barrier, and accumulate in the brain, leading to dementia symptoms [10–12]. There is also evidence suggesting that the gut is a significant source of Aβ in human brains [13]. Taken together, these findings indicate that intestinal bacteria contribute to Aβ accumulation in the brain and, consequently, to the pathophysiology of AD.

In an experimental model, sterile amyloid precursor protein/phosphatidylserine synthase-1 (APP/PSS1) transgenic mice were transplanted with feces from APP/PSS1 transgenic and wild-type mice. Aβ accumulation in the brain was markedly reduced in the group transplanted with feces from wild-type mice compared to the group transplanted with feces from APP/PSS1 mice [14]. This suggests that the intestinal microbiota of wild-type mice may harbor bacteria that inhibit Aβ accumulation.

Based on these findings, we hypothesized that the human intestinal microbiota may also contain bacteria that promote Aβ production (i.e., bacteria contributing to AD pathogenesis) and others that suppress Aβ accumulation (i.e., bacteria mitigating AD progression). Although numerous studies have evaluated the gut microbiota in AD, geographical variations in gut microbiota composition are likely. Notably, relatively little work has been conducted in the Japanese population. Hence, this study was conducted to address the gap in the existing literature. The aim was to investigate the association between microbiota composition and the occurrence of AD in a Japanese population and to clarify whether distinct bacterial species in the human intestinal microbiota are associated with a diagnosis of AD. We successfully identified bacterial species that appeared to be relatively more abundant in patients with AD and others that were more prevalent in healthy adults.

## Materials and methods

### Participants

This exploratory study was conducted at Akirudai Hospital, with sample collection and testing taking place between April 1, 2023 and May 15, 2023. The study included 13 healthy adults aged 22–79 years (mean age: 35.3 years; 7 males and 6 females) and 10 patients with AD aged 78–92 years (mean age: 83.7 years; 2 males and 8 females). Healthy adults were further subdivided into three groups based on age: HV-1 (7 participants aged ≤30 years; 5 males and 2 females), HV-2 (2 participants aged 31–40 years; 1 male and 1 female), and HV-3 (4 participants aged ≥41 years; 1 male and 3 females). Participants provided written informed consent to take part in the study, including for sample collection and testing.

#### Inclusion and Exclusion Criteria

Healthy participants were outpatients without AD who consented to participate, had no known diseases, maintained a balanced diet, and were not taking medications that could influence their intestinal microbiota. There were no additional exclusion criteria for healthy participants. For AD participants, inclusion criteria required a formal diagnosis of AD by a dementia specialist, alongside consent to participate. Exclusion criteria included the use of drugs known to affect intestinal microflora and dietary intolerance.

#### Diagnosis of Alzheimer’s Disease

All AD participants were outpatients from the Neurology Department of Akirudai Hospital. A specialist with 30 years of experience in diagnosing dementia conducted the assessments. The diagnosis followed established AD criteria [15] and included intellectual, cognitive, executive, and memory function tests conducted in a face-to-face format alongside neuropsychological evaluations. Imaging studies, including computed tomography (CT), magnetic resonance imaging (MRI), or amyloid positron emission tomography (PET), were performed to confirm brain and hippocampal atrophy and Aβ accumulation.

#### Dietary and Medication Surveys

Dietary habits, which strongly influence intestinal microbiota composition, were assessed using a brief self-administered diet history questionnaire. Participants were also surveyed regarding the use of oral medications that could potentially impact gut microbiota composition [16].

Fecal sampling was performed after obtaining verbal and written informed consent from each donor, following the ethical principles outlined in the Declaration of Helsinki. The study protocol was approved by the Ethics Review Committee of Akirudai Hospital.

### Analysis of fecal microbiota

Commercially available stool collection kits (Techno Suruga Lab Co. Ltd, Shizuoka, Japan) were used to collect stool samples. The bacterial flora in the collected fecal samples was analyzed using amplicon sequencing, heatmap hierarchical clustering, and diversity analysis (details in S1 Text).

#### Amplicon sequencing analysis

DNA extracted from fecal samples was pretreated according to the method described by Takahashi et al. [17], and the crude DNA was purified using the GENE PREP STAR PI-480 DNA automatic separator (Kurashiki Boseki, Japan). PCR amplification targeted the V3-V4 region of the bacterial 16S rRNA gene using the primer set 341f - R806 [18,19] that amplifies the V3-V4 region of the bacterial 16S rRNA gene according to the method described by Takahashi et al. [17] An index sequence [20] unique to each sample was inserted into each primer at this step. Sequencing was performed using Miseq (Illumina, USA) and MiSeq Reagent Kit v 3 (600 cycles) (Illumina) for paired-end sequencing at 2 × 301 bp cycles.

#### Analysis of the amplicon sequencing data

After paired-end sequencing, primer sequences were removed using Cutadapt version 1.18 [21], and paired-end sequences were joined using Fastq-join software [22] with default settings. Reads in which 99% or more of the bases satisfied a Quality Value (QV) of ≥20 were retained using FASTX-Toolkit version 0.0.14 [23]. Chimeric sequences were removed using usearch61 software [24,25].

Taxonomic classification of nucleotide sequences was performed using Metagenoem@KIN version 2.2.1 software with the RDP version 2.13 database [26] and the microbial identification database NGS-DB-BA version 16.04 [27] (Techno Suruga Lab Co., Ltd., Japan). Taxonomic group confidence was determined using RDP software at ≥0.8, and microbial identification database homology was set at ≥97%.

#### Data analysis using the statistical analysis software R

Using the species-level taxa estimated in the microbial identification database, bacterial species associated with AD and healthy participants were identified. Additionally, the composition of specific taxa, including genera *Bacteroides*, *Blautia*, *Phocaeicola*, the family Lachnospiraceae, and the order Clostridiales, was analyzed. Reversion 3.1.01 software [28] was used to generate boxplots based on grouping information derived from the extracted species composition ratios.

Statistical comparisons were performed using the Mann–Whitney U test [29] with the vegan package version [30,31] in R. The proportions of each bacterial species were evaluated for significant differences among the study groups. To control the false positive rate, P-values were corrected using the Bonferroni method [32]. Statistical significance was set at P<0.05.

## Results

As shown in Figs 1 and 2, some bacterial species appeared more abundant in the AD group, while others were relatively more abundant in healthy adults, with notable variation among the HV subgroups (HV-1, HV-2, and HV-3). However, despite the visible trends in these figures, the differences in bacterial composition among the four groups (AD, HV-1, HV-2, and HV-3) were not statistically significant.

**Fig 1.**
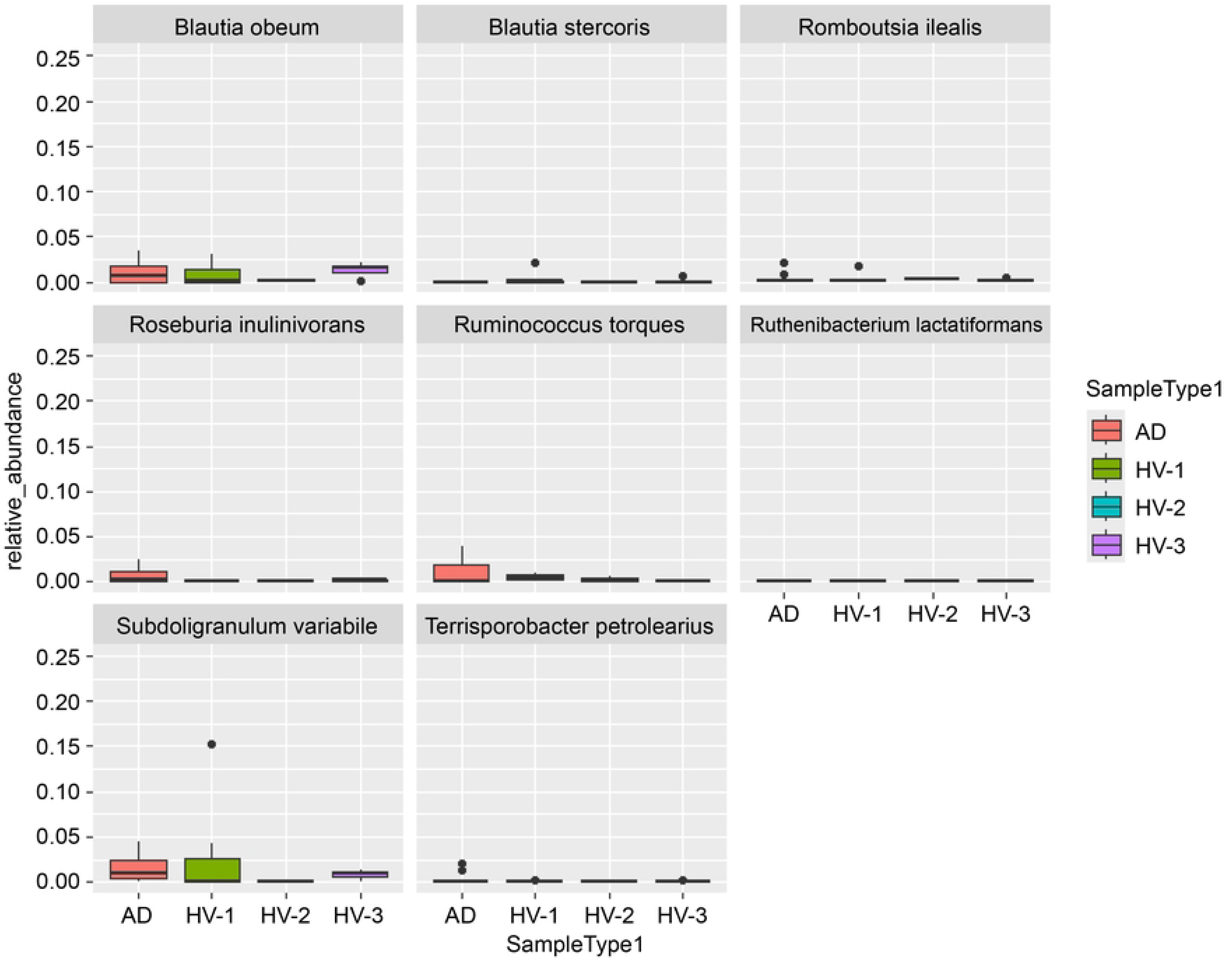
Bacterial species potentially associated with AD. AD, Alzheimer’s disease; HV-1, adults aged ≤30 years; HV-2, adults aged 31–40 years; HV-3, adults aged ≥41 years.

**Fig 2.**
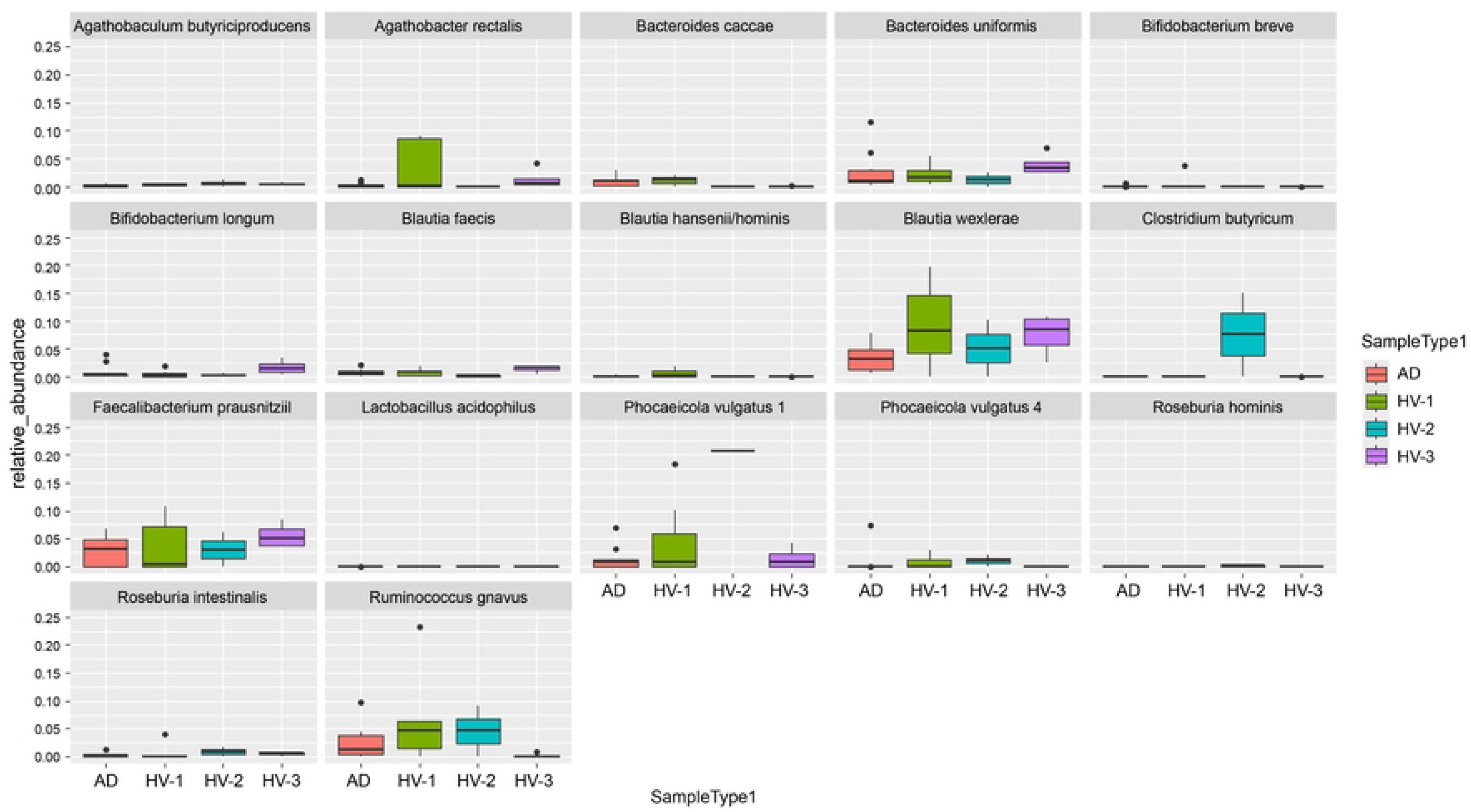
Bacterial species potentially associated with a non-AD state. AD, Alzheimer’s disease; HV-1, adults aged ≤30 years; HV-2, adults aged 31–40 years; HV-3, adults aged ≥41 years.

Among individuals with AD, levels of *R. inulinivorans* and *R. torques* appeared to be non-significantly higher than in healthy individuals (Fig 1). Conversely, levels of *P. vulgatus 1*, *B. wexlerae*, *C. butyricum*, and *A. rectalis* appeared to be higher in healthy individuals, although no significant differences were observed across the different age groups of healthy adults (Fig 2).

The results of the heatmap hierarchical cluster analysis and diversity analysis are provided in S1 Text, alongside S1 and S2 Tables and S1–S8 Figs. Collectively, these analyses indicate that the diversity of bacterial flora differed between the AD and normal subject groups.

## Discussion

The treatment of AD dementia is particularly challenging because of the simultaneous occurrence of neurological damage to brain tissue and mitochondrial dysfunction caused by Aβ accumulation [33]. Lecanemab, a novel monoclonal antibody designed to degrade accumulated Aβ, has been developed as a treatment for AD [34]; another monoclonal antibody—donanemab—has also been evaluated in a phase 2 clinical trial, where it was found to improve cognition and performance in activities of daily living at 76 weeks [35], and has been recently approved for clinical use. While these two treatments could be expected to slow the progression of AD dementia, they are not curative treatments. Indeed, there are currently no curative treatments for AD. Thus, it is essential to screen for mild cognitive impairment (MCI) and AD in their early stages to prevent the progression to AD dementia.

Previous studies have suggested the potential of screening for MCI and early AD by measuring plasma Aβ levels [36,37]. However, no effective screening methods for identifying individuals with pre-AD have been established. In clinical cases of type 2 diabetes mellitus, intestinal bacteria play a key role in maintaining homeostasis until dysbiosis occurs [38], and some authors have suggested the potential to predict the onset of type 2 diabetes mellitus by analyzing the dynamics of intestinal bacteria [39]. Similarly, changes in intestinal microbiota composition and function have been observed to precede the onset of AD, suggesting that analyzing the intestinal microbiota could be a useful method to identify individuals at high risk of developing AD [40]. These changes in intestinal microbiota are detectable prior to alterations in blood markers.

In humans, Aβ begins to accumulate in the brain at approximately 40 years of age, with the onset of MCI and AD occurring roughly 20 years later. During this intermediate period preceding AD onset, cognitive function appears normal [41]. Recent studies using experimental mouse models have demonstrated that intestinal bacteria produce Aβ, which enters the bloodstream, crosses the blood–brain barrier, and accumulates in brain tissue, ultimately leading to dementia [11,12]. Furthermore, emerging evidence suggests that the gut serves as a significant source of Aβ in human brains [13]. While the precise relationship between changes in intestinal microbiota and brain diseases remains unclear, the regulation of dementia pathology via the brain–gut axis has become an increasingly prominent area of research.

In this exploratory study, we identified several species of intestinal bacteria that appeared to be relatively more abundant in the AD group and others that were more prominent in healthy adults. We further investigated the function of these bacteria. Among the species more abundant in the AD group, *R. inulinivorans* is a representative bacterium of the major Clostridial Cluster XIVa group, known for utilizing anaerobic polysaccharides to produce butyric acid during growth. Another species, *R. torques*, is the most efficient degrader of mucin 2, a key component of colonic cell surface mucin. Increased intestinal permeability caused by *R. torques* activity has been implicated in dementia with Lewy bodies, suggesting potential relevance to AD pathogenesis [42]. *R. torques* is also known to inhibit intestinal inflammation by producing secondary bile acids. Butyrate, a short-chain fatty acid produced by certain intestinal bacteria, is generally recognized for promoting mucin production, which strengthens the mucosal layer and supports intestinal barrier function. Additionally, butyrate increases the expression of trefoil factor family peptides, which contribute to the maintenance and repair of intestinal mucosa and enhance the expression of tight junctions in colonic epithelial cells.

However, the effects of butyrate on the intestinal barrier are complex and concentration dependent. While moderate concentrations of butyrate enhance intestinal barrier function, excessively high concentrations can induce apoptosis in colonic epithelial cells, disrupting the intestinal barrier and potentially contributing to colorectal cancer and other diseases. The precise role of butyrate in maintaining intestinal barrier integrity remains incompletely understood and warrants further investigation [43–49]. Reports describing the optimal concentrations of short-chain fatty acids (SCFAs) such as butyric acid, acetic acid, and propionic acid for anti-tumor and anti-inflammatory effects are limited, and further research is required to address this knowledge gap.

Many aspects of bacterial ecology and functions remain poorly understood. Bacteria can incorporate genetic information through crosstalk with their host or environment, exhibiting new functions, including those resulting from horizontal gene transfer (HGT) events. A typical example of HGT is the acquisition of antibiotic resistance by bacteria. In multiple sclerosis (MS), an autoimmune demyelinating disease influenced by environmental factors and sharing some neurodegenerative pathways with AD, HGT has been reported to shape the pathogenicity of certain bacteria [50]. It is plausible that some of the bacterial genetic information mentioned above has been altered because of HGT events.

HGT analysis is not feasible with shotgun metagenomics, but long-read metagenomics enables precise analysis of bacterial chromosomes, making it a valuable tool for studying HGT. Long-read metagenomics can simultaneously detect and visualize HGT events in chromosomes across multiple bacterial species, facilitating detailed functional analysis of bacteria, including plasmid and bacteriophage interactions [51,52].

We hypothesize that the bacterial species potentially associated with AD dementia, as reported in this study, may possess Aβ-generating properties. To explore this hypothesis, we plan to analyze the functional genes of each intestinal bacterium comprehensively using long-read metagenomics, which will allow a more detailed understanding of the genetic mechanisms underlying their potential roles in AD pathogenesis.

We also investigated the functions of intestinal bacteria that were relatively more abundant in healthy adults. *C. butyricum* and *A. rectalis* are butyrate-producing bacteria, contributing to gut barrier integrity and anti-inflammatory effects. *B. wexlerae* produces metabolites such as ornithine, acetylcholine, and S-adenosylmethionine, which inhibit fat accumulation and work cooperatively with other intestinal bacteria to improve the intestinal environment, potentially preventing or mitigating obesity and diabetes. *P. vulgatus 1*, a strain isolated from healthy individuals, exhibits antitoxin, anti-irritable bowel syndrome (IBS), and anti-obesity effects.

Although various bacterial species have demonstrated anti-inflammatory, anti-obesity, and glycolytic effects, only *B. breve* and *A. butyriciproducens* have been reported to prevent dementia. We hypothesize that the bacterial species mentioned above may also exhibit anti-dementia or Aβ-decomposing effects. To investigate this hypothesis, we plan to use long-read sequence metagenomics for comprehensive intestinal microbiota gene analysis.

Analyzing the functional roles of intestinal bacteria must account for the effects of their metabolites. While genome analysis provides insights into bacterial genetic capabilities, metabolome analysis is equally essential for understanding bacterial functions.

Factors influencing intestinal microbiota composition include sex, dietary habits, and medications [53,54]. Among these, dietary habits and oral medications are particularly significant. Multivariate analyses incorporating metadata as confounding factors have revealed that drugs such as gastrointestinal medications, anti-diabetic drugs, antibiotics, antithrombotic agents, cardiovascular and brain disease treatments, and anticancer drugs, in that order, have the most substantial effects on gut microbiota [55]. Specifically, proton pump inhibitors, potassium-channel acid blockers, osmotic laxatives, amino acids, and bile acid inhibitors have strong effects among gastrointestinal drugs, while alpha-glucosidase inhibitors have notable effects among anti-diabetic medications.

Nagata et al. [55] also reported that multi-drug administration increases the number of opportunistic pathogens among commensal bacteria and decreases species producing SCFAs, such as butyric and acetic acid. In our study, although some participants were taking antihypertensive drugs, none were taking medications known to strongly affect gut microbiota or multiple drugs simultaneously. Additionally, none of the participants exhibited dietary deviations.

The results of our exploratory research identified bacterial species that appeared to be relatively more abundant in the AD group and others that were relatively more abundant in the healthy adults. These findings suggest the potential existence of an intermediate group between AD patients and healthy controls. This intermediate group, referred to as the "AD reserve group," may represent individuals at heightened risk of developing dementia. This hypothesis requires validation in future large-scale clinical trials. Based on these findings, various AD prevention projects could be initiated. Currently, we are considering an AD prevention project using the bacterial species identified in this study.

Furthermore, we are exploring the use of artificial intelligence (AI) to analyze the ratio (P/S) of bacterial species associated with AD onset (P) to those associated with AD suppression (S). We hypothesize that the P/S ratio progresses in the following order: healthy group < pre-AD group < MCI/AD group. If validated, this hypothesis could enable rigorous diagnosis of the pre-AD group. This approach has the potential to become a groundbreaking diagnostic tool for pre-AD screening. Moreover, improving the composition of the intestinal microbiota through dementia-protective foods or the transplantation of AD-suppressing bacteria into individuals in the pre-AD group could serve as a preventive strategy against AD [56–59].

Additionally, examining the dynamics of bacteria associated with AD and non-AD states before and after the ingestion of anti-dementia foods and food ingredients could facilitate rigorous evaluation of their cognitive effects. Such research could also enable the development of novel anti-dementia foods and ingredients, further contributing to AD prevention strategies.

There are several limitations to the current study. First, this was a single-center study involving only 23 participants, necessitating repetition of the study with a larger, multi-center sample to validate the findings. Second, the average age of the healthy adult population (35.3 years) was much lower than that of the AD population, which is particularly relevant given that the microbiome changes gradually with age [60]. To address this limitation, we included three groups of healthy adults with different age ranges and analyzed potential age-related differences in microbiota composition. However, the small sample size likely precluded the detection of statistically significant differences, further emphasizing the need for larger studies. Third, while our findings suggest potential associations between specific bacterial species and AD, they do not establish causation.

Based on the analysis of fecal bacteria from healthy individuals and patients with AD, we identified bacterial species in the human intestinal microbiota that could potentially be differentially associated with AD and non-AD states. These findings offer promising insights into the potential for gut microbiota as a modifiable factor in AD prevention. However, large-scale clinical trials and detailed functional analyses of these bacterial species using advanced metagenomic techniques are essential to validate these observations.

Furthermore, given the potential for gut microbiota reconstitution through supplementation [61], long-term interventional studies are needed to assess whether modifying the gut microbiome can influence the risk of developing AD, particularly in at-risk groups. Such studies would contribute significantly to the development of novel prevention strategies and interventions for AD.

## Authors’ contributions

- **Tadashi Ohara**: Conceptualization; Data Curation; Methodology; Formal Analysis; Investigation; Visualization; Writing - Original Draft
- **Yasuyuki Taki**: Supervision; Project Administration
- **Shinya Yamamoto**: Data Curation

## Acknowledgements

The authors extend their gratitude to Dr. Yoichi Imura, president of the medical foundation Akatsuki, Akirudai Hospital, and its staff for their contributions to this study. Technical support from Techno Suruga Lab (https://www.tecsrg.co.jp/) for conducting the sequencing experiments is gratefully acknowledged. Editorial support, in the form of medical writing, assembling tables and creating high-resolution images based on authors’ detailed directions, collating author comments, copyediting, fact checking, and referencing, was provided by Editage, Cactus Communications.

## Data availability statement

The data supporting the findings of this study are available within the article and its supplementary material.l.

---

**S1 Text. Methods and Results for the heatmap hierarchical cluster analysis and diversity analysis**

**S1 Table. Results of pairwise comparisons for alpha diversity**

**S2 Table. Results of pairwise comparisons for beta diversity**

**S1 Fig. Heatmap hierarchical cluster analysis results at the species level**

**S2 Fig. Heatmap hierarchical cluster analysis results at the genus level**

**S3 Fig. Heatmap hierarchical cluster analysis results at the family level**

**S4 Fig. Heatmap hierarchical cluster analysis results at the order level**

**S5 Fig. Heatmap hierarchical cluster analysis results at the class level**

**S6 Fig. Heatmap hierarchical cluster analysis results at the phylum level**

**S7 Fig. Alpha diversity rarefaction plots for the (A) Chao1, (B) Shannon, and (C) Simpson indices at a sequencing depth of 19,049 sequences per sample.** AD: Alzheimer’s disease; HV: healthy adults

**S8 Fig. Three-dimensional principal component analysis plots showing (A) Bray–Curtis distances, (B) unweighted unifrac distances, and (C) weighted unifrac distances visualized using the emperor plot tool function in QIIME2.** The red dots represent the AD group, the blue dots the HV-1 group, the orange dots the HV-2 group, and the green dots the HV-3 group.

## Notes

### Competing Interest Statement

The authors have declared no competing interest.

